# Gene expression is stronger associated with behaviour than with age and fertility in ant workers

**DOI:** 10.1101/267336

**Authors:** Philip Kohlmeier, Austin R Alleman, Romain Libbrecht, Susanne Foitzik, Barbara Feldmeyer

**Affiliations:** Department of Evolutionary Biology, Johannes Gutenberg University Mainz, Johannes von Müller Weg 6, Mainz 55128, Germany; Senckenberg Biodiversity and Climate Research Centre, Senckenberg Gesellschaft für Naturforschung, Senckenberganlage 25, D-60325 Frankfurt am Main, Germany

## Abstract

The ecological success of social insects is based on division of labour, not only between queens and workers, but also among workers. Whether a worker tends the brood or forages is strongly influenced by age, fertility and nutritional status, with brood carers being younger, more fecund and corpulent. Here, we experimentally disentangle behaviour from age and fertility in *Temnothorax longispinosus* ant workers and analyse how these parameters are linked to whole-body gene expression. Our transcriptome analysis reveals four times more genes associated with behaviour than with age and only few fertility-associated genes. Brood carers exhibited an upregulation of genes involved in lipid biosynthesis, whereas foragers invested in metabolism. Additional simulations revealed that the experimental disassociation of co-varying factors reduces transcriptomic noise, potentially explaining discrepancies between transcriptomic studies on worker behaviour in other social insects. Our study highlights the influence of nutritional status on task choice in ant workers.

## Introduction

Division of labour can be found on all levels of biological organization and has arisen independently during several of the major transitions in evolution, for example during the transition from single cell to multicellular organisms or from solitary to social life [1]. The primary driver for the repeated evolution of division of labour is that it allows cells or group members to specialize on few tasks, thus increase in efficiency and allowing for anatomic or physiological adaptations [2–5].

The mechanistic underpinnings regulating specialization are complex and intensively investigated on different levels of biological organisation. A highly developed example of division of labour is found in the societies of ants, bees, wasps and termites. In these colonies of social insects, reproduction is monopolized by a single or very few individuals, the queens (and kings in termites), whereas the majority of the colony members – the workers - remain functionally sterile. All other colony chores are distributed among the workers, which specialize on specific tasks such as brood care, nest defence and foraging [3].

Whether a worker stays inside the nest and cares for the brood or leaves the nest and forages is strongly influenced by age and physiology with brood carers being younger [3], more corpulent [6–14] and more fertile [15,16] than foragers. Due to high external mortality outside the nest [17], this age- and physiology-based non-reproductive division of labour is beneficial, because it ensures that young workers with longest residual life span, the largest amount of stored resources and the highest reproductive potential stay inside the safety of the nest.

Although age and physiology are important regulators of worker task choice in many social insects, few common mechanisms linking age and physiology to behaviour have been found. For example, the switch from brood care to foraging is at least partly regulated by an interaction between age and stable lipid loss [6–14,18,19]. Information on the nutritional status are transported to and in the brain via Target of rapamycin (TOR) and Insulin/Insulin-like signalling (IIS) pathways [12,20–23]. However, transcriptomic patterns and expression biases of multiple behaviour-associated genes seem to be lineage-specific, despite crosstalk with the highly conserved TOR and IIS pathways. Task-specific signatures of nutritional status and metabolism characterize the brood carer- and forager-specific transcriptomes in bees and wasps [18,22,24–29], but such evidence is lacking in ants [30,31]. Brood caring behaviour in honey bees is associated with low titres of juvenile hormone (JH) and high expression of the gene coding for the yolk protein precursor *Vitellogenin* (*Vg*). Once JH titres rise, *Vg* is downregulated resulting in a reduction in brood care, a forager-like gene expression, alterations in small RNA populations, a mobilization of carbohydrates and precocious foraging [26,32–37]. In other social insects, this linkage between JH, *Vg* and behaviour is similar [23,38–40], absent [41] or reversed [42–44]. Moreover, *Vg* underwent multiple duplications in ants with different orthologs taking over specific functions in the regulation of fertility and behaviour [31,45–49]. In honey bees, the *foraging* gene is highly expressed in the optical lobes and mushroom bodies of forager’ brains [50,51], causing an elevated sucrose responsiveness and the onset of foraging behaviour [52,53]. The expression of *foraging* in other social insects is either positively [54–56] or negatively [57–59] linked to foraging behaviour and additionally influenced by age [60], social structure [30], body size [54,58] and time of the day [61]. The manganese transporter *malvolio* is upregulated in foragers, which induces precocious foraging behaviour by increasing sucrose responsiveness in honey bees [62]. Similar to *Vg, malvolio* underwent duplication and subfunctionalization in multiple social and subsocial insects, which raises the question as to whether the role of *malvolio* in honey bees can be extended to other insects [63]. The neuromodulators tyramine and its precursor octopamine promote the onset of foraging, are involved into the rewarding system in honey bee foragers and increase gustatory responsiveness [28,64–66], but their role in other species is unknown.

This across-species inconsistency regarding the mechanistic underpinnings of worker task choice may reflect lineage-specific mechanisms regulating behaviour or differences across studies in the experimental design. Gene expression patterns associated with worker behaviour are typically identified by comparing gene expression of brood carers and foragers. As age and fertility additionally influence gene expression [22,28,67,68], studies that did not experimentally control for age and physiology when comparing gene expression between brood carers and foragers [e.g. 28,31,46,56,57,69–73] might have produced results driven not only by behaviour, but also by age and fertility. Such confounded transcriptomic analyses are not a problem specific to research on insect behaviour but occur across study systems and contexts. For instance, the degree of tissue heterogeneity, i.e. different numbers of cell types present in a tissue, potentially confounds studies comparing gene expression of different tissues [73]. Thus, confounding factors are an important and partly neglected problem when investigating the transcriptomic underpinnings and mechanisms of different traits in general and non-reproductive division of labour in particular.

In this study, we altered the age structure of colonies of the acorn ant *Temnothorax longispinosus* to experimentally create young and old brood carers and foragers respectively. Furthermore, we sampled both fertile and infertile individuals for each combination of behaviour and age. This allowed us to properly assess how behaviour, age, and fertility are associated with gene expression in ant workers. A recent study revealed stronger differences in gene expression between ant queens and workers [68] than between different age-cohorts of the same caste. As we sampled from two clearly distinct age-cohorts differing in age by at least one year, which is estimated to be the mean *T. longispinosus* worker lifespan in the field [74], we expected age to have a stronger impact on gene expression than behaviour. Furthermore, we expected a rather weak association with fertility as we analysed ant workers from queen-right colonies. Nevertheless, a first study on this species revealed differences between fertile and infertile workers [31].

In a second step, we used different subsets of our data to investigate the additional variance introduced to a dataset by not controlling experimentally for confounding factors like age. Transcriptome studies on worker behaviour coupled with such a manipulation have so far been restricted to honey bees [67,75], where they provided great insights into the interaction between behaviour, age and gene expression.

## Material and methods

### Sample collection and preparation

We collected 38 monogynous colonies of the ant *T. longispinosus* with an average colony size of 29 ± 1.5 (mean ± sd) workers at the E.N. Huyck Preserve, Rensselearville, NY, USA in June 2014. Because of the synchronized annual brood emergence in this population around July and August, all field collected workers were at least one year old (termed “old”). *Temnothorax* queens are singly-mated which reduces the potential genetic effects on worker behaviour [76]. In our laboratory in Mainz, Germany, the ant colonies were transferred to slide glass nests and kept in plastered 3-chambered nest-boxes (Figure S1). We maintained them at a 14h:10h light:dark photoperiod and a +22°C:+18°C temperature regime to facilitate brood development. Ants were fed with crickets and honey three times a week. We randomly marked either all young (upon eclosion in the lab) or all old (field collected) workers with thin metal wires (0.02mm, Elektrisola) in each colony, allowing us to differentiate between the two age cohorts. Disentangling the effects of behaviour and age on gene expression requires a full factorial design regarding behaviour (brood carer, forager) and age (young, old). To achieve this, we manipulated the demography of colonies 28 days after the emergence of a new worker generation (Figure 1). Specifically, we either removed (i) all old workers from the colony to induce foraging behaviour in young workers (n=12), (ii) removed all young workers to force old workers to tend the brood (n=10), or (iii) removed half of each age cohort as a control (n=16). Workers were then given 21 days to adjust their behaviour and social organisation. Six brood carers (observed tending the brood) and six foragers (found outside the nest) were then individually labelled with metal wire and their investment into either brood caring and foraging was observed and recorded ten times a day for three days. In *Temnothorax* ants, a single observation is sufficient to classify workers into brood carers and foragers [77] (Kohlmeier et al. subm.). Behavioural observations revealed a clear age-polyethism with young workers focussing on brood care and older workers caring for the adult nestmates and taking over foraging (for details on the methods and results of the behavioural observations see Kohlmeier et al. subm.). After the completion of all observations, we dissected these workers on ice to assess their fertility (fertile: at least one oocyte in the ovaries; infertile: no eggs in development). Following this, individuals were homogenized in 100μl Trizol (Invitrogen) and frozen at −80°C until further processing.

**Figure 1:**
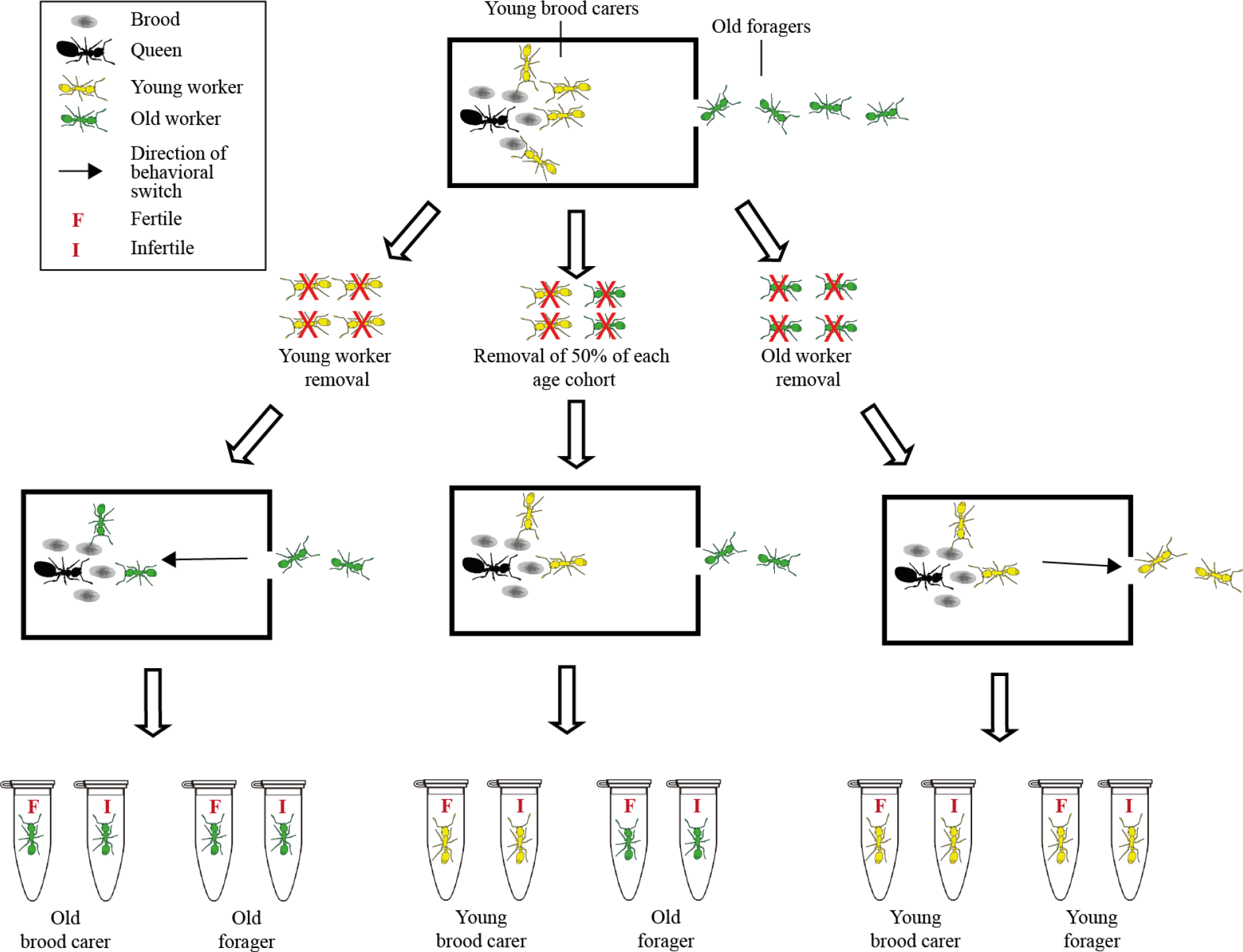
Overview of colony demography manipulations. Colony demographics were manipulated to generate foragers and brood carers of both age classes. From each colony we sampled eight brood carers and eight foragers half of each fertile and half of each unfertile.

Eight brood carers and eight foragers per demography treatment, i.e. a total of 48 workers, were sampled for individual whole-body RNAseq analyses (Figure 1). Muscular activity und consequently behaviour are directly controlled by the brain. However, brain processes are influenced by hormonal activity, nutritional status or ovary development. To gain insights into such up-stream causative factors, we decided to investigate changes in gene expression within the entire worker ant. As *Temnothorax* workers are monomorphic, gene expression differences based on differences in morphology or anatomy are unlikely [78]. As we sampled both, fertile and infertile workers for each combination of behaviour and caste, we were able to investigate gene expression associated with behaviour, age and fertility independent from each other. After defrosting, 100μl Chloroform was added to each sample, after which samples were cautiously inverted for 30s and then centrifuged at 12,000rpm for 15 min at +4°C. The resulting supernatant was collected, and RNA precipitated with 60μl 70% ethanol. Subsequent RNA extraction was conducted with the RNeasy Mini Kit (Qiagen), following the manufacture instructions. Libraries were constructed at GENterprise GmbH Mainz following the standard Illumina protocol, and each library was individually indexed. All 48 libraries were pooled and sequenced with 100 bp paired-end equally spread across eight lanes of an Illumina HiSeq 2500 (Table S1). Sequences will be stored at NCBI short read archive.

### Gene expression analysis

Quality analyses of raw reads were conducted with FastQC (http://www.bioinformatics.babraham.ac.uk/projects/fastqc/) and IIllumina adapters were removed using Trimmomatic [79]. A *de novo* assembly of raw reads was built in two steps: Firstly, remaining raw reads were assembled using CLC Workbench v.7.0.3 (https://www.qiagenbioinformatics.com/; Table S2) and a subsequent meta-assembly was performed in MIRA [80] using CLC Workbench contigs as input. We removed redundant and/or low-confidence contigs from the transcriptome using CD-Hit-Est v.4.6.1 [80].

For the gene expression analysis, reads were aligned to their corresponding contigs using TopHat v2.1.1 [81], in conjunction with Bowtie 2 v2.1.0 (http://bowtie-bio.sourceforge.net/). Read count information was obtained with eXpress v1.5.1 (http://bio.math.berkeley.edu/eXpress/).

Gene expression analysis was performed using *edgeR* v3.6 [82] by running the following three different GLMs to identify genes associated with (i) *behaviour*: brood carers and foragers were compared whereas fertility and age were added as blocking factors, (ii) *age*: old and young workers were compared and fertility and behaviour were added as blocking factors, and (iii) *fertility*: fertile and infertile workers were compared and age and behaviour were added as a blocking factor. The introduction of blocking factors was necessary as samples were not organized in a symmetric full factorial design resulting in, for example, age biases when comparing brood carers to foragers (16 young and 8 old brood carers versus 8 young and 16 old foragers, Figure 1). Contigs that were differentially expressed (FDR < 0.05), but exhibited low expression difference (fold change between groups < 1) were removed. This included 3247 behaviour-, 296 age- and 13 fertility-associated genes. One of the contigs (>philip_contigs_output_c3380) was identified as a chimeric contig containing two similar sized different ORFs (one first ORF was annotated as *Vg3*, the second ORF was annotated as *Vg*1, Kohlmeier et al. subm.). We manually split this contig between both ORFs, remapped and re-counted raw reads and rerun the gene expression analysis. For each of the three factors (behaviour, age, fertility), separate gene onthology (GO) enrichments were performed for both, over- and underexpressed genes, using Blast2GO v4.1.9 [83]. Enriched GO terms were then summarized using ReViGO [84]. A Weighted Gene Coexpression Network Analysis (WGCNA) [85] was performed on the 20,000 contigs exhibiting the strongest variance in expression. Eigengene values were extracted and their association was analysed with a GLMM using *behaviour*, *age*, *fertility* and their interactions as explanatory factors and *colony ID* as a random factor.

We further assessed the expression of candidate genes, which have previously been identified to be involved in the regulation of division of labour in social insects: *foraging*, *Insulin like growth factor*, *Insulin receptor 1* and *2*, *Krueppel homolog 1*, *Tyramine receptor 2*, all orthologs of *Vitellogenin* (*VgC*, *MVg2*, *MVg3*, *Vg-like A*, *Vg-like B*, *Vg-like C*, *Vg receptor*), *ultraspiracle* [18,24,28,86,87]. To identify *foraging* in our transcriptome, we blasted our transcriptome against a database containing 16 foraging genes from different ant species, honey bee and *Drosophila melanogaster*. We only changed the annotation of a specific contig to *foraging*, if the e-value of *foraging* was lower than the e-value of the previous blast result. This resulted in eleven *foraging* contigs.

### Characterizing the effect of age on behaviour specific gene expression

Many studies compared gene expression of young brood carers and old foragers to identify genes associated with behaviour [e.g. 25,28,53,54,66–70]. We investigated the effect of not controlling for age by comparing gene expression of young brood carers (n = 8) and old foragers (n = 8) from control colonies (Figure 1) using pairwise comparison (age confounded PWC) in *edgeR* (e.g. used in [31,88–92]). To test whether differences between age confounded PWC and the age controlled GLM (described in the previous section) depend on whether the dataset was confounded with age or not or on the statistical approach (pairwise comparison *vs*. GLM), we additionally run a GLM on this age confounded dataset including fertility as a blocking factor (age confounded GLM). We compared the number and identity of differentially expressed genes (DEGs) to those, identified with the age controlled GLM. Functional enrichments and WGCNA were performed as described above.

As differences between both age confounded and age controlled approaches can also be explained by differences in sample size (age confounded dataset: 8 brood carers *vs* 8 foragers; age controlled dataset: 24 brood carers *vs* 24 foragers), we randomly sampled four young brood carers, four old brood carers, four young foragers and four old foragers (two of each fertile and two infertile) from our dataset and identified DEGs with GLM including fertility as a blocking factor using 1000 permutations (Reduced sample size GLM; RSS GLM). The total number of possible combinations of samples was 907,200. No combination of samples was used twice.

## Results

### DEGs associated with behaviour independent from age and fertility

A total of 448 genes were differentially expressed between the two behavioural castes (226 overexpressed in brood carers, 222 overexpressed in foragers). 54 of these DEGs were also differentially expressed between young and old workers and 32 between fertile and infertile workers (Figure 2).

**Figure 2:**
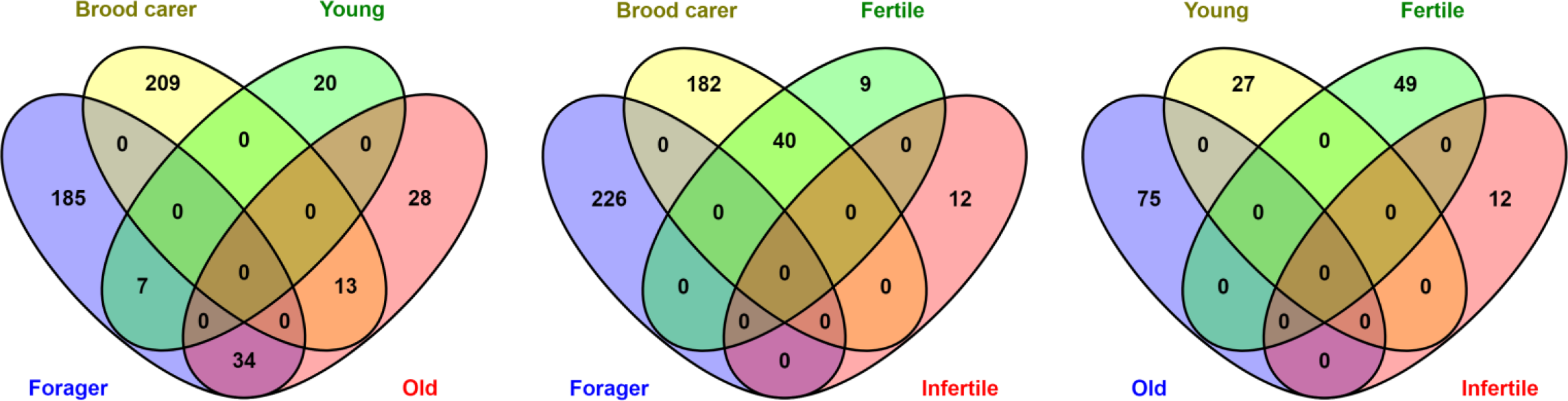
Venn diagrams of genes found to be differentially expressed between brood carers and foragers (left and center) young and old workers (left and right) and fertile and infertile workers (center and right) and their overlap with each other.

Among those genes overexpressed in brood carers, we found several copies of *Vitellogenin* including *Vg-like A, Myrmicine Vg2* (according to Kohlmeier et al. subm.), *MVg3* and the *Vg-Receptor* with *Vg-like A* being the strongest overexpressed gene in brood carers (FDR = 5.45 × 10^−18^). The expression of *VgC, Vg-like B*, and *Vg-like C* was independent from behavioural caste (Table 1). The expression of all differentially expressed *Vg* orthologs was positively correlated with each other (Table S3). Especially the expression of *MVg2* and *MVg3* was tightly linked to each other. Moreover, we found *Neuronal acetylcholine receptor subunit alpha-3*, which binds the neurotransmitter acetylcholine [93], and the *Odorant binding protein 16*, which is part of the olfactory system in honey bees [70]. Enriched GO terms were grouped into eight biological processes, mainly represented by lipid transport and lipid biosynthesis (Figure 3).

**Table 1:**
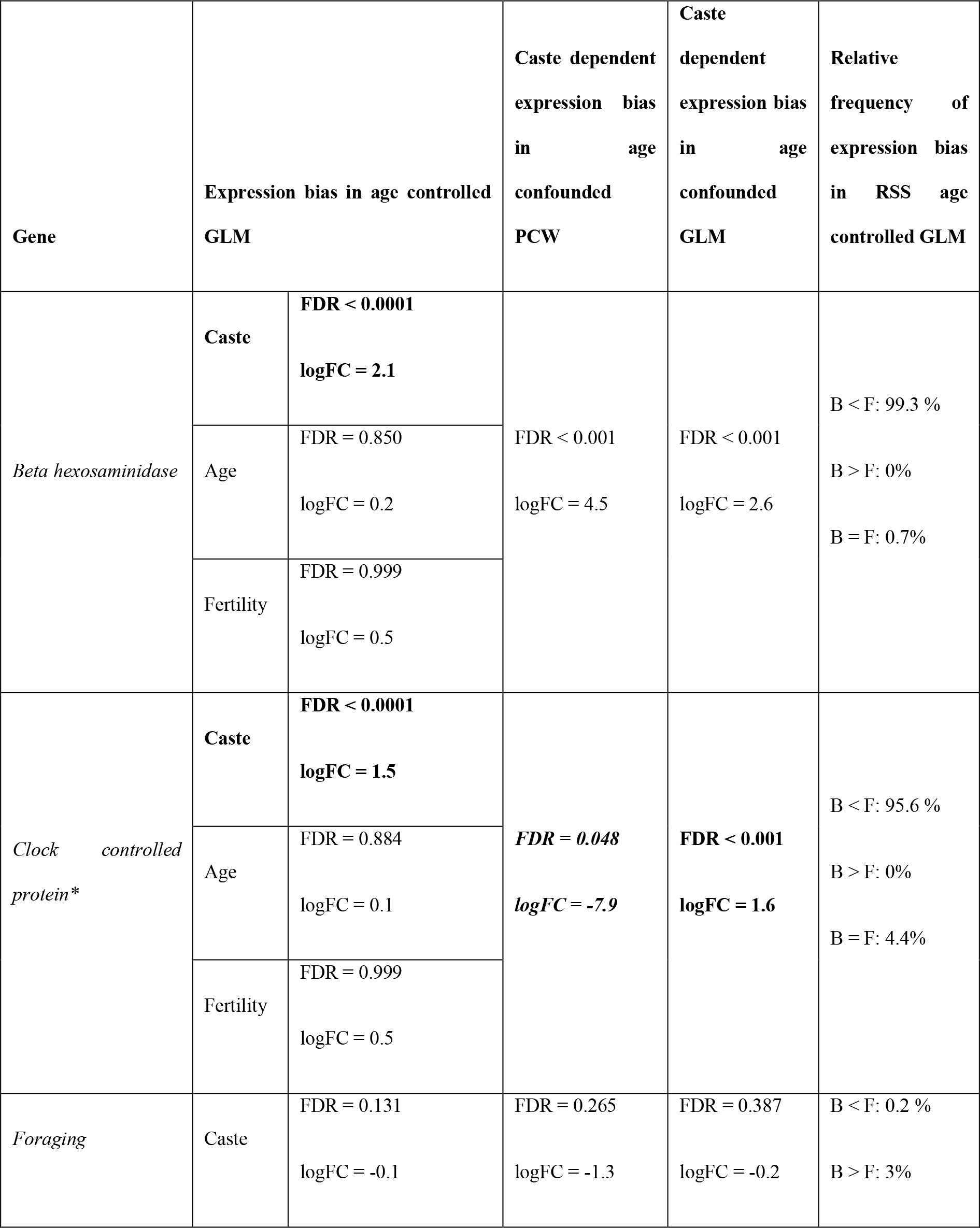
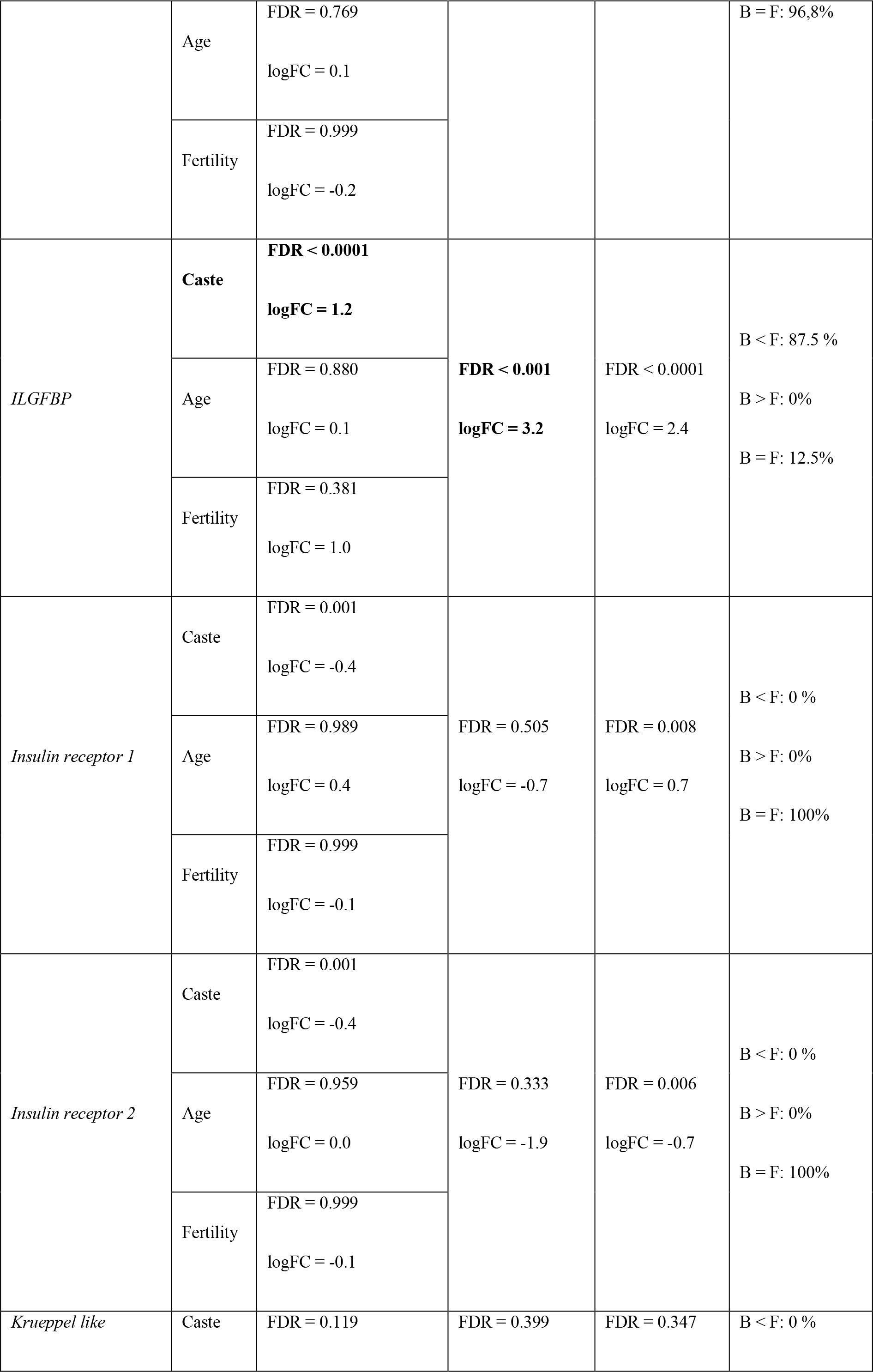
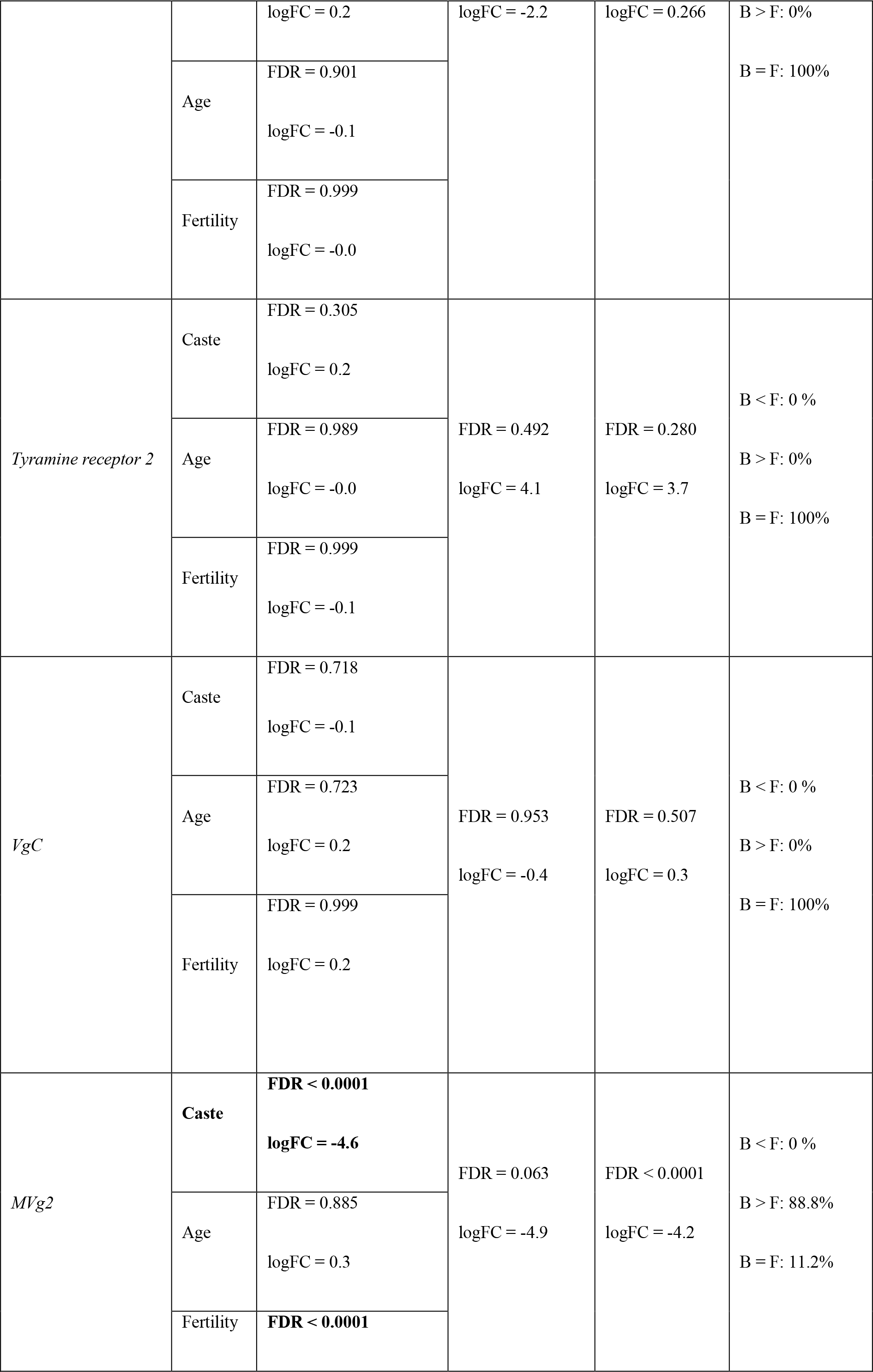
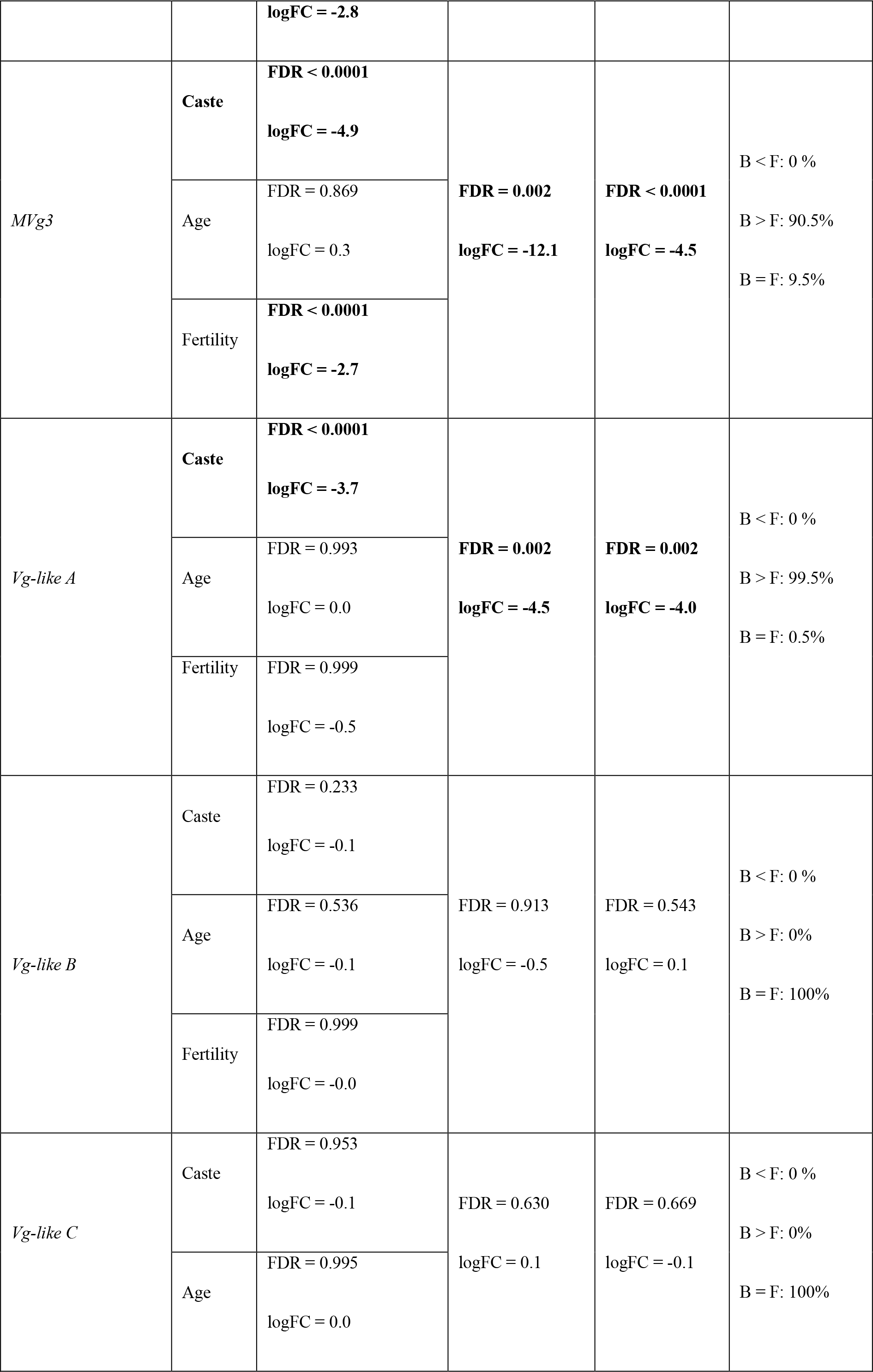
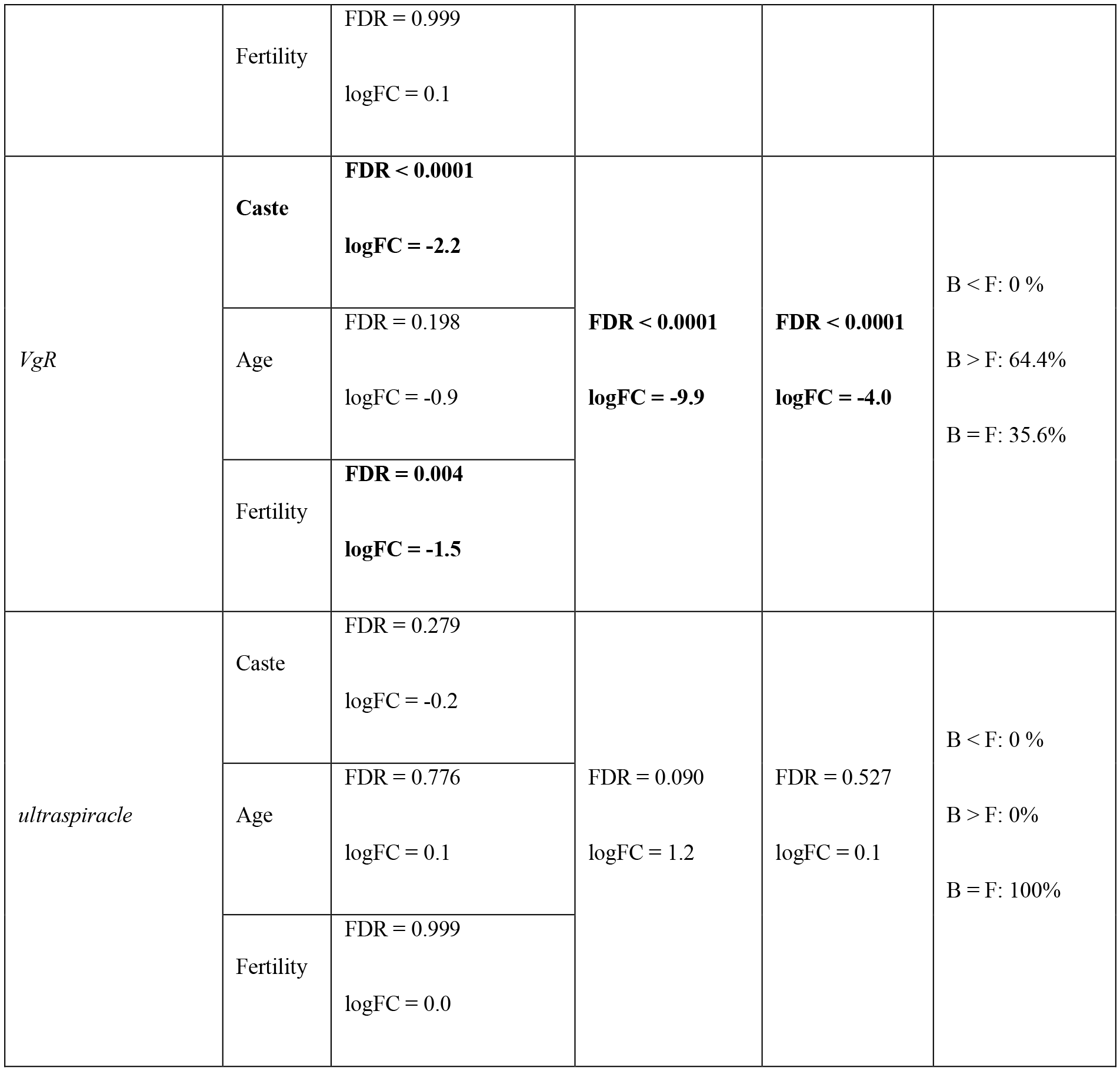
Overview of the candidate genes previously described to be associated with worker behaviour and their expression bias using different methods and datasets. A log fold change (logFC) < 0 represents an expression bias towards brood carers (caste), young (age) or fertile (fertility) workers. Expression was only considered as biased if FDR < 0.05 and logFC < −1 or > 1. Significant expression biases are given in bold. Genes labelled with * exhibited different caste expression biases across the different methods. PCW: Pairwise comparison; RSS: Reduced sample size.

**Figure 3:**
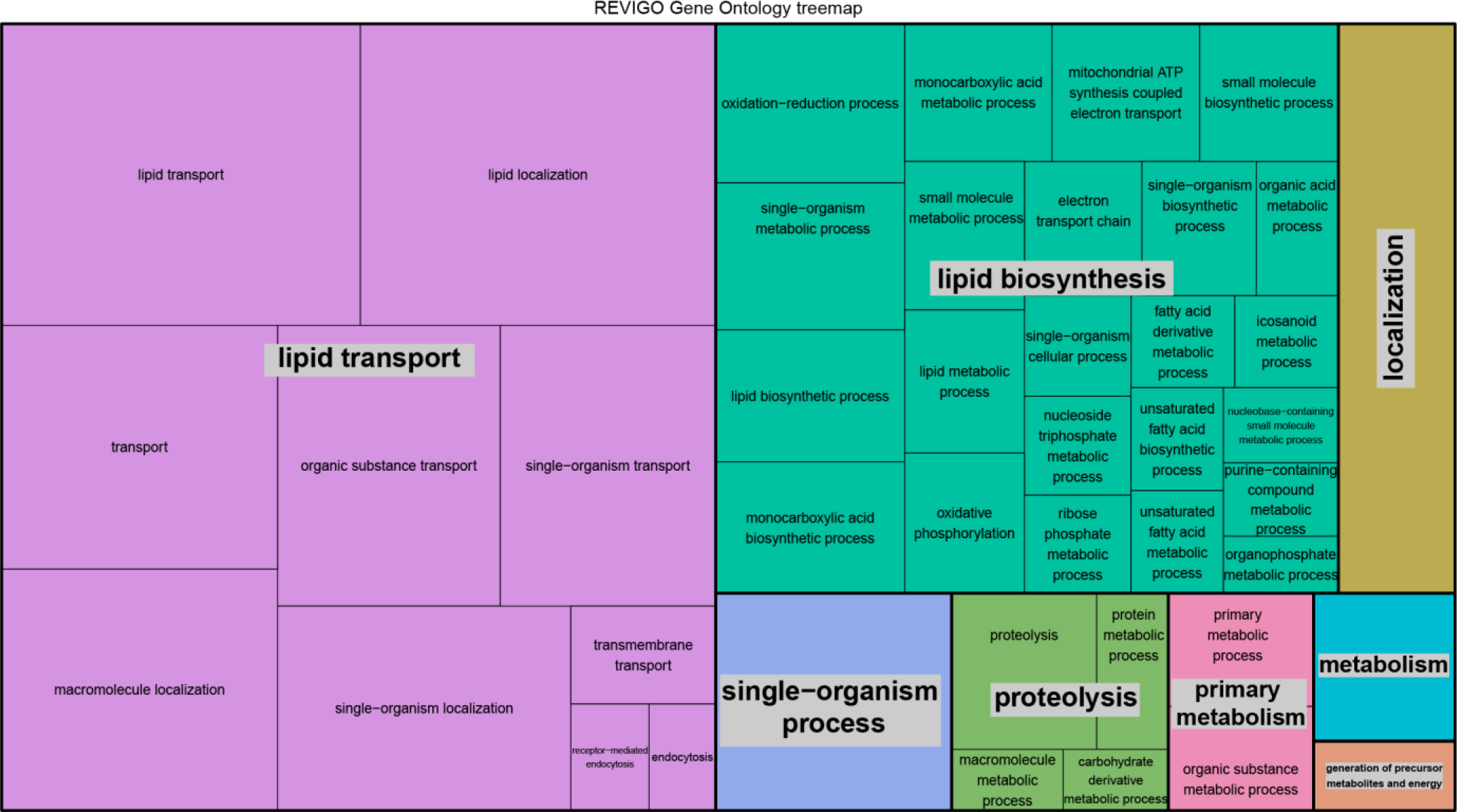
Biological processes overrepresented in the list of genes upregulated in brood carers compared to foragers independent from age and fertility.

Genes overexpressed in foragers include *Insulin-like growth factor-binding protein complex acid labile chain (ILGFBP*), which is associated with binding the Insulin-like growth factor. Furthermore, *beta hexosaminidase subunit beta* and the *circadian clock-controlled protein* were more expressed in foragers than in brood carers. *Tyramine* and *foraging* were not differentially expressed (Table 1). Out of those genes overexpressed in foragers, 20.3% were of viral origin (15.3% *Formica exsecta* virus 2, 2.7% Deformed wing virus, 0.9% *Varroa destructor* virus-1 and Kakugo virus, 0.5% *Nilaparvata lugens* honeydew virus-3). Enriched GO terms were summarized into 13 biological processes largely represented by RNA-templated transcription, pteridine-containing compound catabolism and multiple processes linked to metabolism (Figure 4).

**Figure 4:**
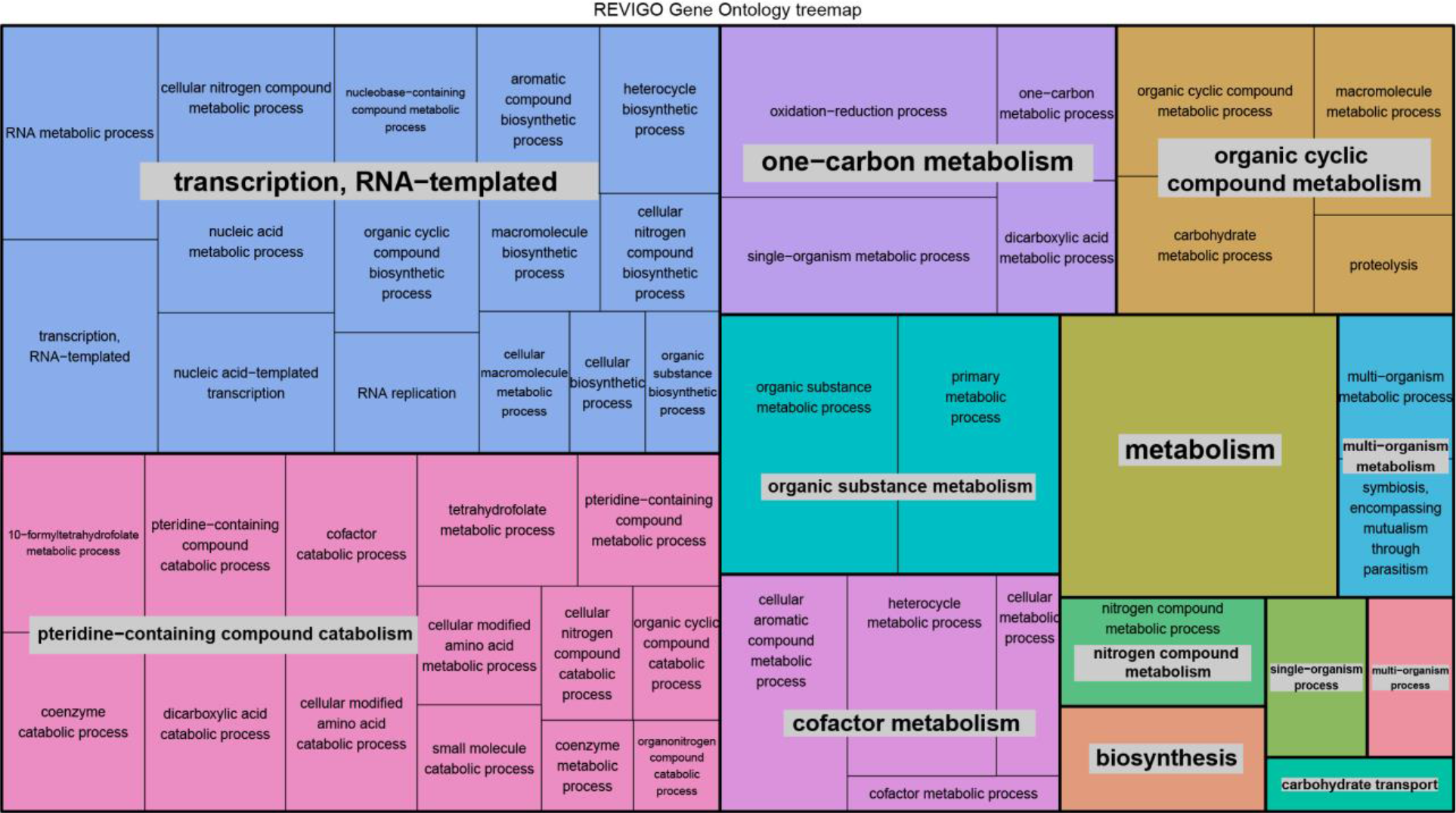
Biological processes overrepresented in the list of genes upregulated in foragers compared to brood carers independent from age and fertility.

Six modules of co-expressed genes were identified using WGCNA (Table S4). One of these modules was associated with brood care behaviour (GLMM: χ^2^ = 4.1 p = 0.042), and exhibited a functional enrichment in metabolism, monocarboxylic acid biosynthesis and biosynthesis.

### DEGs associated with age independent from behaviour and fertility

102 genes were differentially expressed between the two age classes (27 overexpressed in young, 75 in old; Figure 2). The number of these age-specific DEGs is smaller compared to caste-specific DEGs (χ^2^ test: p < 0.0001), but higher than fertility specific DEGs (χ^2^ test: p = 0.001).

Among those genes overexpressed in young workers, we detected multiple *cytochrome P450* genes including *CYP4AB2 and CYP4AB1* that were previously found to exhibit a worker-specific expression in the fire ant *Solenopsis invicta* (Liu and Zhang 2004). Moreover, the expression of *Elongation of very long fatty acids protein*, potentially involved in cuticular hydrocarbon synthesis, *Transposable element P transposase* and multiple copies of the muscle protein *actin*. All genes overexpressed in young workers were combined to a single biological process (single-organism metabolism; Figure S2).

Out of the 75 genes overexpressed in old versus young workers, 36% were of viral origin including *Formica exsecta* virus 2 (28%), Deformed wing virus (4%), Kakugo virus (2.6%) and *Spodoptera frugiperda* rhabdo-virus (1.3%). Ten biological processes including RNA-templated transcription, cellular aromatic compound metabolism, and biosynthesis were linked to old age (Figure S3).

### DEGs associated with fertility independent from behaviour and age

A total of 61 genes were differentially expressed between fertile and infertile workers: 49 of them were overexpressed in fertile and 12 in infertile workers (Figure 2). Among those genes overexpressed in fertile workers, we found *MVg2* (FDR = 3.29 × 10^−5^), *MVg3* (FDR < 0.003) and *Vitellogenin-receptor* (FDR < 0.018). The expression of *VgC*, *Vg-like A*, *Vg-like B* and *Vg-like C* was independent from fertility status (all FDR > 0.999). Six biological processes including lipid transport and mitochondrial electron transport were overrepresented in fertile compared to infertile workers (Figure S4). In infertile workers, the only overrepresented biological process was L-phenylalanine metabolism (Figure S5).

### The combined effect of behaviour and age on gene expression

We used an age confounded subset of the data to characterize the importance of experimentally controlling for worker age. Pairwise comparisons between young brood carers and old foragers only (age confounded PWC) revealed a total of 917 DEGs, significantly more than the number found when controlling for age (χ^2^ test: p < 0.0001; Figure 5a). Out of these, 565 were overexpressed in brood carers and 352 overexpressed in foragers resulting in more overexpressed genes in brood carers than in foragers (χ^2^ test: p < 0.0001), which was not found when controlling for age (χ^2^ test: p = 0.841). In brood carers, the number of biological processes overrepresented was similar to the ones we found when using the age controlled GLM (3 vs. 8; χ^2^ test: p = 0.132; Figure S6), but fewer biological processes where detected among the DEGs of foragers (2 vs. 13; χ^2^ test: p = 0.001, Figure S7).

**Figure 5:**
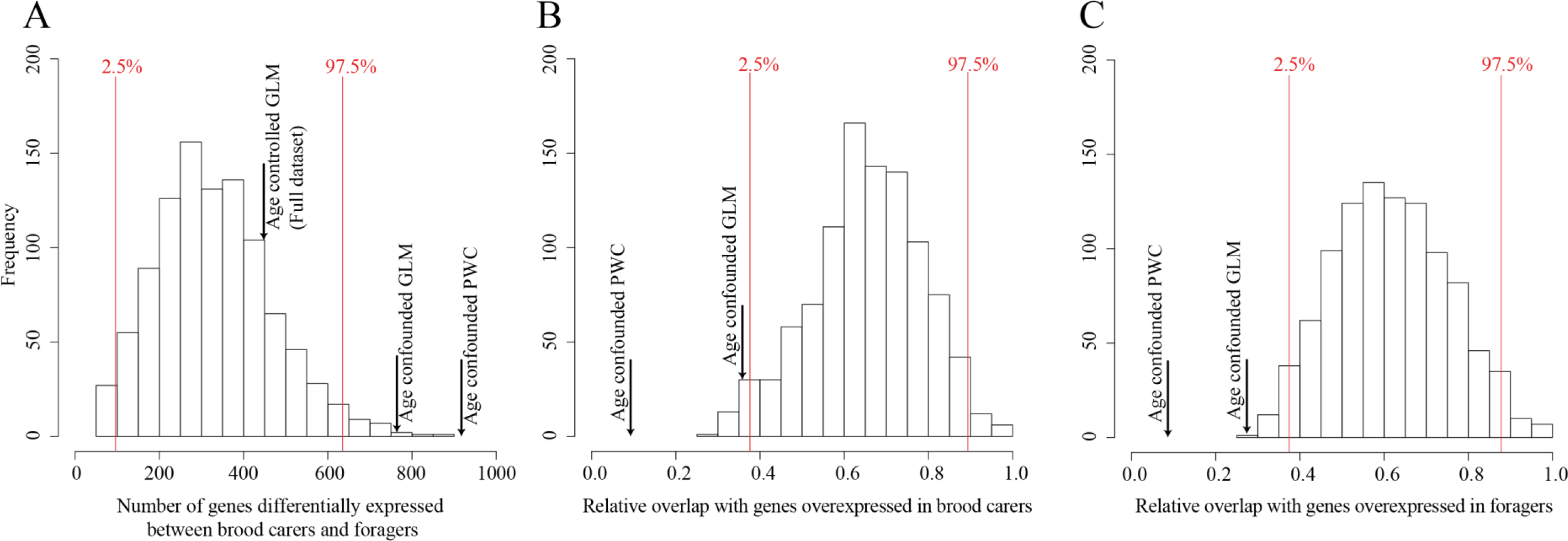
Comparison of age controlled and age confounded gene expression analyses. A) Number of differentially expressed genes identified when comparing brood carers and foragers in an age controlled, reduced sample size GLM (RSS) with 1000 permutations (Bars). B) and C) Relative overlap between the RSS (bars), the age confounded GLM and PWC with the genes found to be upregulated in brood carers (B) and foragers (C) using the age controlled GLM on the full dataset. 5% confidence intervals are given in red. PWC: Pairwise comparison. GLM: Generalized linear model.

The age confounded GLM yielded a total of 764 DEGs between both behavioral castes (Figure 5a): 72.9% more than with the age controlled GLM (χ^2^ test: p < 0.0001), but 16.7% less compared to the age confounded PWC (χ^2^ test: p = 0.0002). When applying a GLM on an age controlled dataset with a reduced sample size (RSS GLM) with 1000 permutations, 330 ± 4.3 DEGs (mean ± S.E.) were identified, which is not different from the 442 DEGS found using the full age controlled dataset (p = 0.813; Figure 5a). The DEGs identified with both age confounded approaches (PWC and GLM) exhibited a lower overlap with those genes identified with the complete age controlled dataset (Brood carer: PWC: 9.2%; GLM: 35.8% figure 5b; Forager: PWC: 8.5%; GLM: 27.3% figure 5c) compared to the age controlled RSS GLM (Brood carer: mean = 65.5%; Forager: mean = 61.7%; figure 5c).

A WGCNA on the age confounded PWC dataset yielded 48 modules. Out of these modules, 14 were associated with brood carers or foragers (Table S5).

## Discussion

In social insects, behavioural specialization of workers (e.g., on brood care and foraging) is influenced by gene expression. Surprisingly, especially in ants, transcriptomic patterns and expression biases of candidate genes involved in the regulation of behaviour vary between studies. The identification of such patterns and genes is difficult, because behaviour is linked to many other factors such as age and fertility. Thus, transcriptome comparisons of brood carers and foragers are often confounded by these parameters, which potentially explains diverging results. In this study, we manipulated colonies of the ant *Temnothorax longispinosus* and assessed gene expression patterns associated with behaviour (brood carer *vs* forager), age (young *vs* old) and fertility (fertile *vs* infertile) independently. In a second step, we compared our results to those that would have been obtained if age was confounded with behaviour.

### Genes associated with behavioural task independent from age and fertility

Our study sheds new light on the transcriptomic underpinnings of non-reproductive division of labour in ants. Gene expression of *T. longispinosus* brood carers was largely represented by lipid transport and lipid biosynthesis (energy storage; Figure 3). Nutritional status, with corpulent brood carers and food deprived foragers, is one of the most widespread and consistent factors mediating worker behaviour [10,19], and influences gene expression in honey bees [12,18,22,25,26,28] and wasps [29]. In ants however, despite the broad correlative evidence for brood carers harbouring more lipid storages than foragers [6–9,14,94,95], clear transcriptomic signatures of nutritional status have so far not been found [30,31]. Only a single study identified lipid storage and fatty acid metabolism to be enriched functions of genes differentially expressed between brood carers and foragers [96]. In that study however, transcriptomes of young brood carers and old foragers were compared. As lipid storages in experimentally hive-restricted bees are reduced with age and age-dependent reduction in lipid storages has been found in *Drosophila* males [12,97], it remained unclear whether observed nutrition linked gene expression is associated with behaviour or with age. Interestingly, we found an increased investment into lipid biosynthesis in brood carers independent from age. In honey bees, reverted brood carers do not refill their lipid storages [19] and future research should therefore test whether old workers in ants, which return to brood caring duties are able to regain lipid storages.

Among the genes upregulated in brood carers, we found multiple fat body expressed copies of egg yolk precursor and storage protein *Vg* (*Vg-like A*, *MVg2*, *MVg3*; Kohlmeier et al. subm.). *Vg-like A* might be of specific importance as its expression was independent from age and fertility, it is mainly expressed in the fat body, has recently been identified as a key regulator of the behavioural transition from brood care to adult nestmate care and occurs in a large number of social insect genomes (Kohlmeier et al. subm.). Whether the changes in responsiveness are at least partly linked to the upregulation of the *Odorant binding protein 16* has to be answered in future. *Conventional Vg* takes over an important role in the regulation of worker behaviour in honey bees [26,32–37]. However, this *Vg* copy was not differentially expressed in our study (see Kohlmeier et al. in subm. for a *Vg* phylogeny). This indicates that brood caring behaviour in ants is controlled by different pathways than in honey bees. Moreover, brood carers overexpressed the brain expressed *Neuronal acetylcholine receptor subunit alpha-3* binds acetylcholine, a neurotransmitter involved in learning in honey bees [93]. However, the role of the receptor for brood caring behaviour should be investigated in the future.

Foragers exhibited increased investment into catabolism and carbohydrate transport (energy mobilization), as well as multiple metabolic processes (energy usage). Similar to honey bees, these data indicate that foragers rely on carbohydrate as a main source of energy for their foraging trips [10]. For example, among the overexpressed genes was *Insulin-like growth factor-binding protein complex acid labile chain*, which is involved in binding of Insulin-like growth factor (IGF) and part of the IIS pathway [98]. Despite the clear transcriptomic signatures of nutritional status and metabolism, multiple candidate genes that are part of the cross-talk between IIS and TOR (i.e. *malvolio, foraging, Tyramine*) were not differentially expressed. As these genes are mainly expressed in the brain [62,99,100], we might have been unable to detect differences in expression, because we used whole body transcriptomes. Alternatively, our findings might reflect lineage specificity regarding pathway rearrangements, which have been documented in the honey bee. For instance, the negative relationship between Insulin-like peptide titres and abdominal lipids in honey bees are a derived stage and reverse in other insects [12,21]. Potentially, such rearrangements have occurred after bee and ant lineages split. This question needs to be answered in the future, e.g. by age controlled tissue-specific gene expression comparisons.

Apart from genes associated with the regulation of behaviour and nutritional physiology, we identified multiple genes that potentially fulfil rather supportive functions for foraging behaviour. Mutations in *Beta hexosaminidase subunit beta*, one of the strongest differentially expressed genes between brood carers and foragers, are linked to the Sandhoff disease in humans causing complex symptoms including a reduced locomotive activity [101]. *Clock-controlled protein* is a gene downstream of the circadian clock in *Drosophila melanogaster* [102]. Whether these genes for example contribute to maintaining muscle activity (*beta hexosaminidase subunit beta*) or correctly timing foraging trips (*clock-controlled protein*) still needs to be investigated.

### Genes associated with age independent from behavioural task and fertility

Young age in workers was characterized by the overexpression of several cytochrome *P450* genes including *CYP4AB 1* and *2*, *Elongation of very long chain fatty acids protein*, *transposable element P transposase* and several actin genes. *P450 CYP4AB1* and *2* were shown to be overexpressed in workers compared to sexuals in the fire ant *Solenopsis invicta* [103]. Overexpression of *Elongation of very long chain fatty acids protein* potentially contributes to age-dependent differences in cuticular hydrocarbon profiles, which have been reported for workers of the ant *Diacamma ceylonese* [104]. Old workers in contrast overexpress viral transcripts, indicative of a higher viral load in aged workers, which was previously documented in honey bees, in which a sugar-rich diet further increased the viral load of workers [105]. High viral load might contribute to the increased intrinsic mortality of old compared to young workers as well as foragers compared to brood carers [77] Interestingly, we did not detect any typical aging pathways or genes, such as ROS pathways.

Age had a much weaker influence on the transcriptome compared to behaviour. This is in line with honey bee age controlled gene expression studies [67,75], although the age difference in honey bee brood carers and foragers is few weeks only, where it is at least one year in our study. Our findings strongly contrast with studies investigating caste (queen *vs*. worker) gene expression across different developmental stages and ages, in which more DEGs were found between different ages or developmental stages than between castes [68,91,106–108]. The larger gene expression differences between brood carers and foragers in combination with no upregulation of pathways associated with longevity indicate strong physiological differences between the two behavioural worker castes, and suggest weak changes with age in the investment in body repair mechanisms [109].

### Genes associated with fertility independent from behavioural task and age

Fertility had the weakest effect on overall gene expression. Noteworthy however, *MVg2* and *MVg3* were highly expressed in fertile compared to infertile workers, whereas the expression of *Vg-like A* was independent from fertility status. This supports the hypothesis of a sub-functionalization of *Vg* and *Vg-like* genes. In future, it will be interesting to see whether the Myrmicine-specific *MVg2* and *MVg3* fulfil fertility-linked functions similar to the ancestral *Vg* copies e.g. in *Drosophila*.

### The combined effect of behaviour and age on gene expression

We created age confounded subdatasets to compare gene expression between young brood carers and old foragers using pairwise comparison (Age confounded PCW) and a GLM including *fertility* as a blocking factor (Age confounded GLM). This allowed us to investigate the effect of not experimentally controlling for age. Despite few exceptions (*MVg2, clock-controlled protein*), candidate genes involved in the regulation of worker behaviour showed similar expression patterns across all methods (Table 1). Thus, investigations of specific candidate genes seem to be consistent even with confounded data and highlight the potential importance of these genes for the regulation of worker behaviour.

However, additional analyses, such as GO enrichment, WGCNA and metabolic pathway comparisons are commonly used and provide valuable insights into broader patterns of gene expression [27,30,31,67,68,75,87,90,96]. Hence, results and conclusions depend on the number and identity of large sets of genes, and partly their FDR values and raw read counts provided as an input. We could show that both age confounded approaches (PWC and GLM) identified more DEGs than the age controlled approaches (RSS and GLM). These findings are likely to be the result of additional variation in gene expression that was introduced by workers not only differing in behaviour but also in age. This additional variation was not a simple combination of those genes found to be differentially expressed between behaviours and age (Figure S8), but seems to be rather random with unpredictable effects on follow-up analyses: Whereas GO term enrichments on the age confounded dataset yielded less enriched biological processes than the age controlled dataset, more co-expressed modules were found in the WGCNA. Thus, our data highlight the necessity of experimentally controlling for confounding factors.

The use of age confounded datasets might therefore have contributed to the lack of consensus on clear transcriptomic patterns of worker behaviour, as many studies investigating gene expression in different behavioural castes did not control for age and fertility [e.g. 28,31,56,57,69–73]. Similar conclusions were recently drawn from a gene expression comparison between queens and workers, as uncontrolled variation was added by confounding age when comparing gene expression between both castes [68]. Such effects of experimentally introduced variation have not only been described for age but also for tissue choice [68,71,72,78]. We therefore conclude that gene expression analyses aiming at identifying genes associated with worker behaviour benefits from experimentally controlling for confounding factors such as age.

### Conclusions

Taken together, by disentangling gene expression associated with behaviour, age and fertility, we identified new candidate genes involved in the regulation of non-reproductive behaviour (*Vg-like A*), showed that candidate genes described in honey bees do not play a role in *Temnothorax* ants (*Conventional Vg*), and highlight the importance of nutritional physiology and metabolism in brood carers versus foragers. In respect to expression changes with age, we detected investment in muscles in young workers, and no evidence for age-dependant expression of typical longevity genes. We thus provide evidence that experimentally controlling for confounding factors such as age or behaviour will increase resolution and decrease transcriptomic noise in future RNAseq studies.

## References

1. Maynard Smith J, Szathmáry E. The major transitions in evolution. Oxford: Oxford University Press; 1995.

2. Smith A. The wealth of nations. UK: Methuen; 1776.

3. Wilson EO. The Insect Societies. Cambridge: Harvard University Press; 1971.

4. Robinson GE. Regulation of division of labor in insect societies. Annu Rev Entomol. 1992;37: 637–665.

5. Wahl LM. Evolving the division of labour: Generalists, specialists and task allocation. J Theor Biol. 2002;219: 371–388. doi:10.1006/jtbi.2002.3133

6. Porter SD, Jorgensen CD. Foragers of the harvester ant, Pogonomyrmex owyheei: a disposable caste? Behav Ecol Sociobiol. 1981;9: 247–256. doi:10.1007/BF00299879

7. Tschinkel WR. Seasonal life history and nest architecture of a winter-active ant, Prenolepis imparis. Insectes Soc. 1987;34: 143–164. doi:10.1007/BF02224081

8. Tschinkel W. Sociometry and sociogenesis of the harvester ant Pogonomyrmex badius: worker characteristics in relation to colony size and season. Insectes Soc. 1998;45: 385–410.

9. Schulz DJ, Huang ZY, Robinson GE. Effects of colony food shortage on behavioral development in honey bees. Behav Ecol Sociobiol. 1998;42: 295–303. doi:10.1007/s002650050442

10. Toth AL, Robinson GE. Worker nutrition and division of labour in honeybees. Anim Behav. 2005;69: 427–435. doi:10.1016/j.anbehav.2004.03.017

11. Toth AL, Bilof KBJ, Henshaw MT, Hunt JH, Robinson GE. Lipid stores, ovary development, and brain gene expression in Polistes metricus females. Insectes Soc. 2009;56: 77–84. doi:10.1007/s00040-008-1041-2

12. Ament SA, Wang Y, Robinson GE. Nutritional regulation of division of labor in honey bees: Toward a systems biology perspective. Wiley Interdiscip Rev Syst Biol Med. 2010;2: 566–576. doi:10.1002/wsbm.73

13. Robinson EJH, Feinerman O, Franks NR. Flexible task allocation and the organization of work in ants. Proc R Soc B Biol Sci. 2009;276: 4373–4380.

14. Bernadou A, Busch J, Heinze J. Diversity in identity: behavioral flexibility, dominance, and age polyethism in a clonal ant. Behav Ecol Sociobiol. 2015;69: 1365–1375. doi:10.1007/s00265-015-1950-9

15. Hölldobler B, Carlin NF. Colony founding, queen control and worker reproduction in the ant Aphaenogaster (=Novomessor) cockerelli (Hymenoptera: Formicidae). Psyche (Stuttg). 1989;96: 131–151.

16. Kühbandner S, Modlmeier AP, Foitzik S. Age and ovarian development are related to worker personality and task allocation in the ant Leptothorax acervorum. 2014;60: 392–400.

17. Visscher PK, Dukas R. Survivorship of foraging honey bees. Insectes Soc. 1997;44: 15. doi:10.1007/s000400050017

18. Ament SA, Chan QW, Wheeler MM, Nixon SE, Johnson SP, Rodriguez-Zas SL, et al. Mechanisms of stable lipid loss in a social insect. J Exp Biol. 2011;214: 3808–3821. doi:10.1242/jeb.060244

19. Toth AL, Kantarovich S, Meisel AF, Robinson GE. Nutritional status influences socially regulated foraging ontogeny in honey bees. J Exp Biol. 2005;208: 4641–4649. doi:10.1242/jeb.01956

20. Maestro JL, Cobo J, Bellés X. Target of rapamycin (TOR) mediates the transduction of nutritional signals into juvenile hormone production. J Biol Chem. 2009;284: 5506–5513. doi:10.1074/jbc.M807042200

21. Morton GJ, Cummings DE, Baskin DG, Barsh GS, Schwartz MW. Central nervous system control of food intake and body weight. Nature. 2006;443: 289–295. doi:10.1038/nature05026

22. Nilsen K-A, Ihle KE, Frederick K, Fondrk MK, Smedal B, Hartfelder K, et al. Insulin like peptide genes in honey bee fat body respond differently to manipulation of social behavioral physiology. J Exp Biol. 2011;214: 1488–1497. doi:10.1242/jeb.050393

23. Rodrigues MA, Flatt T. Endocrine uncoupling of the trade-off between reproduction and somatic maintenance in eusocial insects. Curr Opin Insect Sci. Elsevier Inc; 2016;16: 1–8. doi:10.1016/j.cois.2016.04.013

24. Ament SA, Corona M, Pollock HS, Robinson GE. Insulin signaling is involved in the regulation of worker division of labor in honey bee colonies. Proc Natl Acad Sci. 2008;105: 4226–4231. doi:10.1073/pnas.0800630105

25. Zayed A, Robinson GE. Understanding the Relationship Between Brain Gene Expression and Social Behavior: Lessons from the Honey Bee. Annu Rev Genet. 2012;46: 591–615. doi:10.1146/annurev-genet-110711-155517

26. Wang Y, Brent CS, Fennern E, Amdam G V. Gustatory perception and fat body energy metabolism are jointly affected by vitellogenin and juvenile hormone in honey bees. PLoS Genet. 2012;8. doi:10.1371/journal.pgen.1002779

27. Khamis AM, Hamilton AR, Medvedeva YA, Alam T, Alam I, Essack M, et al. Insights into the transcriptional architecture of behavioral plasticity in the honey bee Apis mellifera. Sci Rep. Nature Publishing Group; 2015;5: 11136. doi:10.1038/srep11136

28. Scheiner R, Entler B V., Barron AB, Scholl C, Thamm M. The effects of fat body tyramine level on gustatory responsiveness of honeybees (Apis mellifera) differ between behavioral castes. Front Syst Neurosci. 2017;11: 1–8. doi:10.3389/fnsys.2017.00055

29. Daugherty THF, Toth AL, Robinson GE. Nutrition and division of labor: Effects on foraging and brain gene expression in the paper wasp Polistes metricus. Mol Ecol. 2011;20: 5337–5347. doi:10.1111/j.1365-294X.2011.05344.x

30. Manfredini F, Lucas C, Nicolas M, Keller L, Shoemaker D, Grozinger CM. Molecular and social regulation of worker division of labour in fire ants. Mol Ecol. 2014;23: 660–72. doi:10.1111/mec.12626

31. Feldmeyer B, Elsner D, Foitzik S. Gene expression patterns associated with caste and reproductive status in ants: worker-specific genes are more derived than queen-specific ones. Mol Ecol. 2014;23: 151–161. doi:10.1111/mec.12490

32. Marco Antonio DS, Guidugli-Lazzarini KR, do Nascimento AM, Simões ZLP, Hartfelder K. RNAi-mediated silencing of vitellogenin gene function turns honeybee (Apis mellifera) workers into extremely precocious foragers. Naturwissenschaften. 2008;95: 953–961. doi:10.1007/s00114-008-0413-9

33. Amdam G V, Norberg K, Hagen A, Omholt SW. Social exploitation of vitellogenin. Proc Natl Acad Sci U S A. 2003;100: 1799–802. doi:10.1073/pnas.0333979100

34. Nunes FMF, Ihle KE, Mutti NS, Simões ZLP, Amdam G V. The gene vitellogenin affects microRNA regulation in honey bee (Apis mellifera) fat body and brain. J Exp Biol. 2013;216: 3724–32. doi:10.1242/jeb.089243

35. Guidugli KR, Nascimento AM, Amdam G V, Barchuk AR, Omholt S, Simões ZLP, et al. Vitellogenin regulates hormonal dynamics in the worker caste of a eusocial insect. FEBS Lett. 2005;579: 4961–4965. doi:10.1016/j.febslet.2005.07.085

36. Amdam G V, Norberg K, Page RE, Erber J, Scheiner R. Downregulation of vitellogenin gene activity increases the gustatory responsiveness of honey bee workers (Apis mellifera). Behav Brain Res. 2006;169: 201–205. doi:10.1016/j.bbr.2006.01.006

37. Wheeler MM, Ament SA, Rodriguez-Zas SL, Southey B, Robinson GE. Diet and endocrine effects on behavioral maturation-related gene expression in the pars intercerebralis of the honey bee brain. J Exp Biol. 2015;218: 4005–4014. doi:10.1242/jeb.119420

38. Robinson GE, Vargo EL. Juvenile hormone in adult eusocial hymenoptera: Gonadotropin and behavioral pacemaker. Arch Insect Biochem Physiol. 1997;35: 559–583.

39. Engel KC, Stökl J, Schweizer R, Vogel H, Ayasse M, Ruther J, et al. A hormone-related female anti-aphrodisiac signals temporary infertility and causes sexual abstinence to synchronize parental care. Nat Commun. 2016;7: 1–10. doi:10.1038/ncomms11035

40. Azevedo DO, de Paula SO, Zanuncio JC, Martinez LC, Serrão JE. Juvenile hormone downregulates vitellogenin production in Ectatomma tuberculatum (Hymenoptera: Formicidae) sterile workers. J Exp Biol. 2015;219: 103–108. doi:10.1242/jeb.127712

41. Amsalem E, Malka O, Grozinger C, Hefetz A. Exploring the role of juvenile hormone and vitellogenin in reproduction and social behavior in bumble bees. BMC Evol Biol. BMC Evolutionary Biology; 2014;14: 45. doi:10.1186/1471-2148-14-45

42. Brent CS, Vargo EL. Changes in juvenile hormone biosynthetic rate and whole body content in maturing virgin queens of Solenopsis invicta. J Insect Physiol. 2003;49: 967–974. doi:10.1016/S0022-1910(03)00166-5

43. Libbrecht R, Corona M, Wende F, Azevedo DO, Serrão JE, Keller L. Interplay between insulin signaling, juvenile hormone, and vitellogenin regulates maternal effects on polyphenism in ants. Proc Natl Acad Sci U S A. 2013;110: 11050–5. doi:10.1073/pnas.1221781110

44. Pamminger T, Treanor D, Hughes WOH. Pleiotropic effects of juvenile hormone in ant queens and the escape from the reproduction-immunocompetence trade-off. Proc Biol Sci. 2016;283: 20152409-. doi:10.1098/rspb.2015.2409

45. Wurm Y, Wang J, Riba-Grognuz O, Corona M, Nygaard S, Hunt BG, et al. The genome of the fire ant Solenopsis invicta. Proc Natl Acad Sci. 2011;108: 5679–5684. doi:10.1073/pnas.1009690108

46. Corona M, Libbrecht R, Wurm Y, Riba-grognuz O, Studer RA, Keller L. Vitellogenin underwent subfunctionalization to acquire caste and behavioral specific expression in the harvester ant Pogonomyrmex barbatus. PLoS Genet. 2013;9: e1003730. doi:10.1371/journal.pgen.1003730

47. Oxley PR, Ji L, Fetter-Pruneda I, McKenzie SK, Li C, Hu H, et al. The genome of the clonal raider ant Cerapachys biroi. Curr Biol. 2014;24: 451–458. doi:10.1016/j.cub.2014.01.018

48. Morandin C, Havukainen H, Kulmuni J, Dhaygude K, Trontti K, Helanterä H. Not only for egg yolk-functional and evolutionary insights from expression, selection, and structural analyses of formica ant vitellogenins. Mol Biol Evol. 2014;31: 2181–2193. doi:10.1093/molbev/msu171

49. Salmela H, Sundström L. Vitellogenin in inflammation and immunity in social insects. Inflamm Cell Signal. 2017;4: e1506. doi:10.14800/ics.1506

50. Ben-Shahar Y, Robichon A, Sokolowski MB, Robinson GE. Influence of gene action across different time scales on behavior. Science. 2002;296: 741–744. doi:10.1126/science.1069911

51. Heylen K, Gobin B, Billen J, Hu TT, Arckens L, Huybrechts R. Amfor expression in the honeybee brain: A trigger mechanism for nurse-forager transition. J Insect Physiol. 2008;54: 1400–1403. doi:10.1016/j.jinsphys.2008.07.015

52. Thamm M, Scheiner R. PKG in honey bees: Spatial expression, Amfor gene expression, sucrose responsiveness, and division of labor. J Comp Neurol. 2014;522: 1786–1799.

53. Scheiner R, Page RE, Erber J. The effects of genotype, foraging role, and sucrose responsiveness on the tactile learning performance of honey bees (Apis mellifera L.). Neurobiol Learn Mem. 2001;76: 138–150. doi:10.1006/nlme.2000.3996

54. Tobback J, Mommaerts V, Vandersmissen HP, Smagghe G, Huybrechts R. Age- and task-dependent foraging gene expression in the bumblebee Bombus terrestris. Arch Insect Biochem Physiol. 2011;76: 30–42. doi:10.1002/arch.20401

55. Toth AL, Varala K, Henshaw MT, Rodriguez-Zas SL, Hudson ME, Robinson GE. Brain transcriptomic analysis in paper wasps identifies genes associated with behaviour across social insect lineages. Proc R Soc B Biol Sci. 2010;277: 2139–2148. doi:10.1098/rspb.2010.0090

56. Bockoven AA, Coates CJ, Eubanks MD. Colony-level behavioral variation correlates with differences in expression of the foraging gene in red imported fire ants. Mol Ecol. 2017;12: 3218–3221. doi:10.1111/mec.14347

57. Ingram KK, Oefner P, Gordon DM. Task-specific expression of the foraging gene in harvester ants. Mol Ecol. 2005;14: 813–818. doi:10.1111/j.1365-294X.2005.02450.x

58. Kodaira Y, Ohtsuki H, Yokoyama J, Kawata M. Size-dependent foraging gene expression and behavioral caste differentiation in Bombus ignitus. BMC Res Notes. 2009;2: 1–5. doi:10.1186/1756-0500-2-184

59. Lucas C, Sokolowski MB. Molecular basis for changes in behavioral state in ant social behaviors. Proc Natl Acad Sci U S A. 2009;106: 6351635–6. doi:10.1073/pnas.0809463106

60. Oettler J, Nachtigal AL, Schrader L. Expression of the foraging gene is associated with age polyethism, not task preference, in the ant Cardiocondyla obscurior. PLoS One. 2015;10: e0144699. doi:10.1371/journal.pone.0144699

61. Ingram KK, Kleeman L, Peteru S. Differential regulation of the foraging gene associated with task behaviors in harvester ants. BMC Ecol. BioMed Central Ltd; 2011;11: 19. doi:10.1186/1472-6785-11-19

62. Ben-Shahar Y, Dudek NL, Robinson GE. Phenotypic deconstruction reveals involvement of manganese transporter malvolio in honey bee division of labor. J Exp Biol. 2004;207: 3281–3288. doi:10.1242/jeb.01151

63. Mehlferber EC, Benowitz KM, Roy-Zokan EM, McKinney EC, Cunningham CB, Moore AJ. Duplication and sub/neofunctionalization of malvolio, an insect homolog of Nramp, in the subsocial beetle Nicrophorus vespilloides. Genes|Genomes|Genetics. 2017;7: 3393–3403. doi:10.1534/g3.117.300183

64. Schulz DJ, Barron AB, Robinson GE. A role for octopamine in honey bee division of labor. Brain Behav Evol. 2002;60: 350–359. doi:10.1159/000067788

65. Scheiner R, Plückhahn S, Öney B, Blenau W, Erber J. Behavioural pharmacology of octopamine, tyramine and dopamine in honey bees. Behav Brain Res. 2002;136: 545–553.

66. Barron AB, Maleszka R, Meer RK Vander, Robinson GE. Octopamine modulates honey bee dance behavior. Proc Natl Acad Sci. 2007;104: 1703–1707.

67. Whitfield CW, Cziko A-M, Robinson GE. Gene expression profiles in the brain predict behavior in individual honey bees. Science. 2003;302: 296–299. doi:10.1126/science.1086807

68. Lucas ER, Romiguier J, Keller L. Gene expression is more strongly influenced by age than caste in the ant Lasius niger. Mol Ecol. 2017;38: 42–49. doi:10.1111/mec.14256

69. Cardoso-Júnior CAM, Silva RP, Borges NA, de Carvalho WJ, Walter SL, Simões ZLP, et al. Methyl farnesoate epoxidase (mfe) gene expression and juvenile hormone titers in the life cycle of a highly eusocial stingless bee, Melipona scutellaris. J Insect Physiol. Elsevier; 2017;101: 185–194. doi:10.1016/j.jinsphys.2017.08.001

70. Zhao H, Luo Y, Lee J, Zhang X, Liang Q, Zeng X. The odorant-bindingprotein gene obp11 shows different spatiotemporal roles in the olfactory system of Apis mellifera ligustica and Apis cerana cerana. Sociobiology. 2013;60: 429–435. doi:10.13102/sociobiology.v60i4.429-435

71. Johnson BR, Jasper WC. Complex patterns of differential expression in candidate master regulatory genes for social behavior in honey bees. Behav Ecol Sociobiol. Behavioral Ecology and Sociobiology; 2016;70: 1033–1043. doi:10.1007/s00265-016-2071-9

72. Johnson BR, Atallah J, Plachetzki DC. The importance of tissue specificity for RNA-seq: Highlighting the errors of composite structure extractions. BMC Genomics. 2013;14. doi:10.1186/1471-2164-14-586

73. Erkkilä T, Lehmusvaara S, Ruusuvuori P, Visakorpi T, Shmulevich I, Lähdesmäki H. Probabilistic analysis of gene expression measurements from heterogeneous tissues. Bioinformatics. 2010;26: 2571–2577. doi:10.1093/bioinformatics/btq406

74. Foitzik S, Herbers JM. Colony structure of a slavemaking ant. II. Frequency of slave raids and impact on the host population. Evolution (N Y). 2001;55: 316–323. doi:10.1111/j.0014-3820.2001.tb01296.x

75. Alaux C, Le Conte Y, Adams HA, Rodriguez-Zas S, Grozinger CM, Sinha S, et al. Regulation of brain gene expression in honey bees by brood pheromone. Genes, Brain Behav. 2009;8: 309–319. doi:10.1111/j.1601-183X.2009.00480.x

76. Foitzik S, Herbers JM. Colony structure of a slavemaking ant. I. Intracolony relatedness, worker reproduction, and polydomy. Evolution (N Y). 2001;55: 307–315. doi:10.1111/j.0014-3820.2001.tb01295.x

77. Kohlmeier P, Negroni MA, Kever M, Emmling S, Stypa H, Feldmeyer B, et al. Intrinsic worker mortality depends on behavioral caste and the queens’ presence in a social insect. Sci Nat. The Science of Nature; 2017;104: 34. doi:10.1007/s00114-017-1452-x

78. Weitekamp CA, Libbrecht R, Keller L. Genetics and evolution of social behavior in insects. Annu Rev Genet. 2017;51: 219–239. doi:10.1146/annurev-genet-120116-024515

79. Bolger AM, Lohse M, Usadel B. Trimmomatic: A flexible trimmer for Illumina sequence data. Bioinformatics. 2014;30: 2114–2120. doi:10.1093/bioinformatics/btu170

80. Li W, Godzik A. Cd-hit: A fast program for clustering and comparing large sets of protein or nucleotide sequences. Bioinformatics. 2006;22: 1658–1659. doi:10.1093/bioinformatics/btl158

81. Trapnell C, Pachter L, Salzberg SL. TopHat: Discovering splice junctions with RNA-Seq. Bioinformatics. 2009;25: 1105–1111. doi:10.1093/bioinformatics/btp120

82. Robinson MD, McCarthy DJ, Smyth GK. edgeR: A Bioconductor package for differential expression analysis of digital gene expression data. Bioinformatics. 2009;26: 139–140. doi:10.1093/bioinformatics/btp616

83. Götz S, Garcia-Gómez JM, Terol J, Williams TD, Nagaraj SH, Nueda MJ, et al. High-throughput functional annotation and data mining with the Blast2GO suite. Nucleic Acids Res. 2008;36: 3420–3435. doi:10.1093/nar/gkn176

84. Supek F, Bošnjak M, Škunca N, Šmuc T. Revigo summarizes and visualizes long lists of gene ontology terms. PLoS One. 2011;6. doi:10.1371/journal.pone.0021800

85. Langfelder P, Horvath S. WGCNA: an R package for weighted correlation network analysis. BMC Bioinformatics. 2008;9: 559. doi:10.1186/1471-2105-9-559

86. Grozinger CM, Sharabash NM, Whitfield CW, Robinson GE. Pheromone-mediated gene expression in the honey bee brain. Proc Natl Acad Sci. 2003;100: 14519–14525. doi:10.1073/pnas.2335884100

87. Ament SA, Wang Y, Chen CC, Blatti CA, Hong F, Liang ZS, et al. The transcription factor ultraspiracle influences honey bee social behavior and behavior-related gene expression. PLoS Genet. 2012;8. doi:10.1371/journal.pgen.1002596

88. Landis GN, Abdueva D, Skvortsov D, Yang J, Rabin BE, Carrick J, et al. Similar gene expression patterns characterize aging and oxidative stress in Drosophila melanogaster. Proc Natl Acad Sci. 2004;101: 7663–7668. doi:10.1073/pnas.0307605101

89. Grozinger CM, Fan Y, Hoover SER, Winston ML. Genome-wide analysis reveals differences in brain gene expression patterns associated with caste and reproductive status in honey bees (Apis mellifera). Mol Ecol. 2007;16: 4837–4848. doi:10.1111/j.1365-294X.2007.03545.x

90. Ferreira PG, Patalano S, Chauhan R, Ffrench-Constant R, Gabaldón T, Guigó R, et al. Transcriptome analyses of primitively eusocial wasps reveal novel insights into the evolution of sociality and the origin of alternative phenotypes. Genome Biol. 2013;14. doi:10.1186/gb-2013-14-2-r20

91. Harrison MC, Hammond RL, Mallon EB. Reproductive workers show queenlike gene expression in an intermediately eusocial insect, the buff-tailed bumble bee Bombus terrestris. Mol Ecol. 2015;24: 3043–3063. doi:10.1111/mec.13215

92. Feldmeyer B, Mazur J, Beros S, Lerp H, Binder H, Foitzik S. Gene expression patterns underlying parasite-induced alterations in host behaviour and life history. Mol Ecol. 2016;25: 648–660. doi:10.1111/mec.13498

93. Shapira M, Thompson CK, Soreq H, Robinson GE. Changes in neuronal acetylcholinesterase gene expression and division of labor in honey bee colonies. J Mol Neurosci. 2001;17: 1–12. doi:10.1385/JMN:17:1:1

94. Robinson EJH, Feinerman O, Franks NR. Flexible task allocation and the organization of work in ants. 2015;276: 4373–4380.

95. Silberman RE, Gordon D, Ingram KK. Nutrient stores predict task behaviors in diverse ant species. Insectes Soc. Springer International Publishing; 2016;63: 299–307. doi:10.1007/s00040-016-0469-z

96. Qiu H, Zhao C, He Y. On the molecular basis of division of labor in Solenopsis invicta (Hymenoptera: Formicidae) workers: RNA-seq Analysis. J Insect Sci. 2017;17: 1–9. doi:10.1093/jisesa/iex002

97. Vermeulen CJ, Van De Zande L, Bijlsma R. Developmental and age-specific effects of selection on divergent virgin life span on fat content and starvation resistance in Drosophila melanogaster. J Insect Physiol. 2006;52: 910–919. doi:10.1016/j.jinsphys.2006.05.014

98. Leong SR, Baxter RC, Camerato T, Dai J, Wood WI. Structure and functional expression of the acid-labile subunit of the insulin-like growth factor-binding protein complex. Mol Endocrinol. 1992;6: 870–876.

99. Ben-Shahar Y, Leung H-T, Pak WL, Sokolowksi MB, Robinson GE. cGMP-dependent changes in phototaxis: a possible role for the foraging gene in honey bee division of labor. J Exp Biol. 2003;206: 2507–2515. doi:10.1242/jeb.00442

100. Wagener-Hulme C, Kuehn JC, Schulz DJ, Robinson GE. Biogenic amines and division of labor in honey bee colonies. J Comp Physiol - A Sensory, Neural, Behav Physiol. 1999;184: 471–479. doi:10.1007/s003590050347

101. Sandhoff K, Andreae U, Jatzkewitz H. Deficient hexosaminidase activity in an exceptional case of Tay-Sachs disease with additional storage of kidney globoside in visceral organs. Life Sci. 1968;7: 283–288. Available: http://www.ncbi.nlm.nih.gov/pubmed/5651108

102. Lorenz LJ, Hall J, Rosbash M. Expression of a Drosophila messenger-RNA is under circadian clock control during pupation. Development. 1989;107: 869–880.

103. Liu N, Zhang L. CYP4AB1, CYP4AB2, and Gp-9 gene overexpression associated with workers of the red imported fire ant, Solenopsis invicta Buren. Gene. 2004;327: 81–87. doi:10.1016/j.gene.2003.11.002

104. Cuvillier-Hot V, Cobb M, Malosse C, Peeters C. Sex, age and ovarian activity affect cuticular hydrocarbons in Diacamma ceylonense, a queenless ant. J Insect Physiol. 2001;47: 485–493. doi:10.1016/S0022-1910(00)00137-2

105. DeGrandi-Hoffman G, Chen Y, Huang E, Huang MH. The effect of diet on protein concentration, hypopharyngeal gland development and virus load in worker honey bees (Apis mellifera L.). J Insect Physiol. Elsevier Ltd; 2010;56: 1184–1191. doi:10.1016/j.jinsphys.2010.03.017

106. Hoffman EA, Goodisman MA. Gene expression and the evolution of phenotypic diversity in social wasps. BMC Biol. 2007;5: 23. Available: http://www.ncbi.nlm.nih.gov/entrez/query.fcgi?cmd=Retrieve&db=PubMed&dopt=Citation&list_uids=17504526

107. Ometto L, Shoemaker D, Ross KG, Keller L. Evolution of gene expression in fire ants: The effects of developmental stage, caste, and species. Mol Biol Evol. 2011;28: 1381–1392. doi:10.1093/molbev/msq322

108. Smith CR, Helms Cahan S, Kemena C, Brady SG, Yang W, Bornberg-Bauer E, et al. How do genomes create novel phenotypes? Insights from the loss of the worker caste in ant social parasites. Mol Biol Evol. 2015;32: 2919–2931. doi:10.1093/molbev/msv165

109. Kramer BH, Schaible R. Life span evolution in eusocial workers-A theoretical approach to understanding the effects of extrinsic mortality in a hierarchical system. PLoS One. 2013;8: e61813. doi:10.1371/journal.pone.0061813

